# Inducing synthetic lethality for selective targeting of acute myeloid leukemia cells harboring *STAG2* mutations

**DOI:** 10.1101/2022.02.18.479480

**Authors:** Agatheeswaran Subramaniam, Carl Sandén, Larissa Moura-Castro, Kristijonas Žemaitis, Ludwig Schmiderer, Alexandra Bäckström, Elin Arvidsson, Simon Hultmark, Natsumi Miharada, Kajsa Paulsson, Thoas Fioretos, Jonas Larsson

## Abstract

Targeted therapies exploiting selective vulnerabilities of malignant cells are highly desired for clinical applications. The cohesin protein complex comprises of RAD21, SMC3, SMC1A as well as a fourth subunit that consists of either STAG1 or STAG2 and is essential for proper chromosomal segregation during mitosis. *STAG2* loss-of-function mutations are recurrent driver events in acute myeloid leukemia (AML) and appear relatively early during leukemogenesis. Studies in cell lines have shown that *STAG2* deficient cells are uniquely vulnerable to STAG1 perturbation, and this vulnerability could thus be exploited to selectively eliminate *STAG2* null AML cells. Here we show that partial perturbation of STAG1 is well tolerated by normal human hematopoietic stem cells and does not affect their functionality. By contrast, STAG1 knockdown is lethal to *STAG2* null human HSCs by inducing major mitotic defects. Moreover, STAG1 knockdown induced synthetic lethality in primary human AML cells harboring a *STAG2* mutation and completely abrogated leukemia progression in xenograft models. Overall, our study provides proof-of-concept demonstration of a synthetic lethal approach to selectively target primary human cancer cells with *STAG2* mutations

## Introduction

Acute myeloid leukemia (AML) is a heterogeneous disorder and mostly incurable due to relapse and drug resistance. A key challenge is to target the fraction of more quiescent leukemic cells that are resistant to chemotherapeutic drugs (1). During recent years, several new small molecule inhibitors have been developed to uniquely target disease-specific molecular events, such as *FLT3-ITD* mutation. However, such targeted therapies have only met moderate clinical success due to drug resistance and relapse, which is associated with selection and expansion of malignant subclones that are not dependent on the initially targeted mutation (2). Thus, additional novel targeted therapeutic strategies are required for leukemia clearance. A particularly attractive approach is to harness specific genetic deficiencies of the tumor cells by targeting synthetic lethal interactions. In this case, a therapeutic effect can be reached even in tumors that are not critically dependent on the mutations that are targeted.

The cohesin protein complex forms a ring-like structure that holds sister chromatids together which is necessary for the proper segregation of chromosomes during mitosis. In addition, cohesin plays an essential role in DNA repair, genome organization and transcriptional regulation (3). The core structure comprises the three proteins RAD21, SMC3 and SMC1A. The fourth subunit consists of either one of two paralogous proteins: STAG1 or STAG2. Whole genome sequencing studies have identified a significant number of somatic mutations in cohesin genes with an accumulated mutation rate between 10 and 15 % for AML and MDS (4). An even higher rate of cohesin mutations (around 50%) was observed in Down Syndrome associated childhood acute megakaryocytic leukemia (DS-AMKL) (5). Cohesin mutations have loss-of-function consequences arguing for a tumor suppressor role of cohesin in the context of leukemia (6). Since cohesin is necessary for proper chromosomal segregation, mutations were first thought to promote tumor progression via genome instability (7). However, the majority of the cohesin mutated cancers are euploid suggesting that it is rather non-mitotic functions of cohesin such as transcriptional regulation and chromatin organization which are associated with leukemogenesis (8). With a mutation frequency of approximately 6% in AML and 18% in DS-AMKL, *STAG2* is the predominantly mutated gene among the cohesin genes (4). *STAG2* mutations result in complete loss-of-function in males since these genes are located on the X chromosome (9). STAG2 knockdown promoted the *in vitro* expansion of umbilical cord blood (UCB)-derived hematopoietic stem and progenitor cells (HSPCs) and enhanced the repopulating activity of human HSPCs in xenograft recipients, demonstrating a direct functional association between STAG2 loss and dysregulated hematopoiesis (10). Although STAG1- and STAG2-containing cohesin complexes might have distinct functions (11), they are redundant to ensure the chromatid cohesion during mitosis. Hence, STAG1-mediated mitotic dependency was observed in *STAG2* knockout cell lines (12, 13). Moreover, a recent study in knockout mice showed that the combined loss of STAG1 and STAG2, but not that of each gene alone, resulted in bone marrow aplasia and mortality, further supporting the existence of synthetic lethal interactions between STAG1 and STAG2 (14). A majority of the cohesin mutations have a high variant allele frequency indicating that they occur relatively early in leukemogenesis (4, 15). Inducing synthetic lethality in AML harboring *STAG2* mutations should thus eliminate most clones irrespective of any secondary mutations. Though, theoretically possible as a novel therapeutic application to *STAG2* null AML, synthetic lethality from targeting STAG1 is yet to be demonstrated in primary human leukemic cells. Here, we provide a functional proof-of-concept demonstration for this approach.

## Results and discussion

First, to demonstrate the existence of STAG1 mediated mitotic dependency in primary human HSPCs, we generated *STAG2* null HSPCs derived from UCB utilizing a CRISPR/Cas9 based knock-in system. In brief, an early stop codon, followed by an open reading frame encoding enhanced green fluorescent protein (eGFP) was inserted into the targeted STAG2 locus utilizing homology-directed repair (HDR) (**Supplementary Figure 1A**). With this system, successfully edited, eGFP expressing STAG2 null cells can be distinguished from unedited cells. As STAG2 is located in the X chromosome, we edited CD34^+^ cells from male donors to achieve successful *STAG2* knockout by monoallelic editing. Flow cytometry analysis revealed an eGFP frequency of 27% compared to mock control indicating efficient *STAG2* editing and eGFP integration in HSPCs (**Figure 1A & B**). Further, Western blotting analysis of the eGFP expressing cells revealed a near complete loss of STAG2 protein, demonstrating that integration of the eGFP template successfully knocked out *STAG2* expression (**Figure 1C**). Sanger sequencing of the *STAG2* locus in eGFP positive cells reveled a successful integration of the donor template at the expected locus (Supplementary Figure 1B). Altogether we successfully generated *STAG2* knockout (KO) human HSPCs utilizing CRISPR mediated HDR.

**Figure 1.**
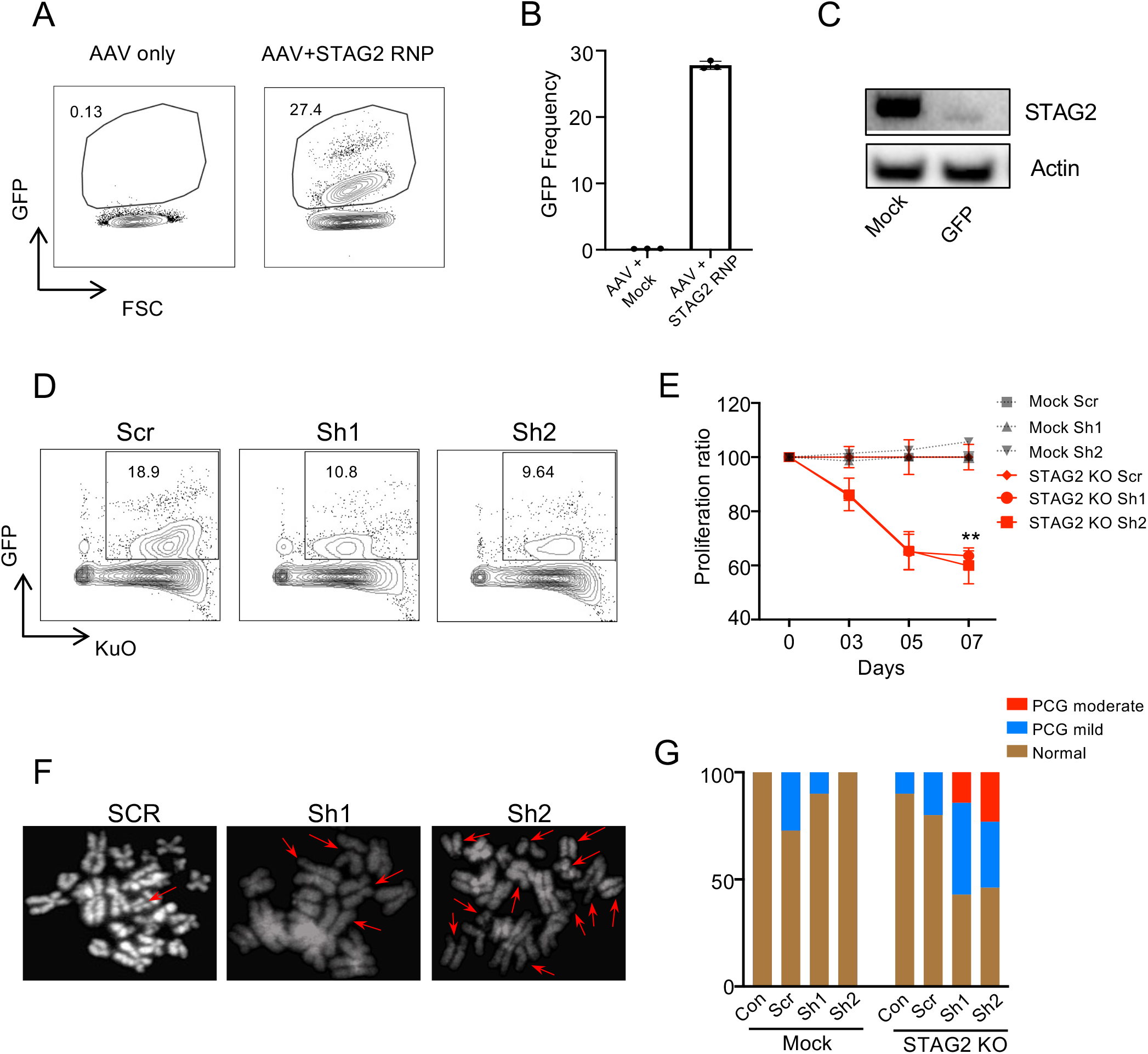
STAG1 knockdown perturbs STAG2 null HSPCs. Umbilical cord blood (UCB) CD34^+^ cells were cultured in serum free expansion medium (SFEM) added with stem cell factor (SCF), thrombopoietin (TPO) and FMS-like tyrosine kinase 3 ligand (FLT3L) at final concentration of 100 ng/mL each. A. Flow cytometry analysis of enhanced green fluorescent protein (eGFP) expression in UCB CD34^+^ cells edited with either mock or with sgRNA (UCUGGUCCAAACCGAAUGAA) - Cas9 ribonucleoproteins targeting STAG2 along with adeno associated virus donor template. B. Quantification of CRISPR/Cas9 mediated eGFP knock-in efficiency measurement across three replicates. C. Western blot analysis of STAG2 protein in the mock and eGFP sorted UCB CD34^+^ cells. D. Day 05 co-transduction analysis of Kusabira orange positive STAG1 shRNAs and eGFP positive *STAG2* null cells by flow cytometry. E. Quantification of STAG1 shRNA mediated cell proliferation in mock and *STAG2* null CD34^+^ cells compared to the scrambled control. Two-way ANOVA, p<0.01 - ***. F. Fluorescence i*n situ* hybridization to analyze the sister chromatid cohesin in *STAG2* null cells 3 days after shRNA transduction. G. Cohesion defects were quantified in around 8-15 cells for each condition. Primary constriction gaps (PCG) measured are the visible gaps between the sister chromatids at the centromeres; PCG mild - defects in 1-4 Chromosomes, PCG moderate - defects in 4-19 chromosomes. Scr-Scrambled shRNA CAACAAGATGAAGAGCACCAA; Sh1-STAG1 shRNA1 CTTCAGCCTTTGGTGTTCAAT; Sh2-STAG1 shRNA2 GCCAATGAAAGGTTGGAGTTA.

Next to assess STAG1 mediated mitotic dependency, we transduced *STAG2* KO human HSPCs with two independent short hairpin RNAs (shRNA) targeting STAG1 as well scrambled control shRNA. The pLKO shRNA vectors were engineered to express Kusabira orange (KuO) enabling tracking of the transduced cells in conjunction with the eGFP marker for *STAG2* null cells. Both STAG1 shRNAs showed successful knockdown of STAG1 at the mRNA and protein level 72 hours post-transduction (**Supplementary figure 1C & D**). We monitored cell number and frequency of the Kusabira orange and GFP positive population during one week of culture and observed that both STAG1 shRNAs, but not the control shRNA, induced a significant depletion of *STAG2* null HSPCs. Moreover, isogenic control cells with intact STAG2 were unaffected by the STAG1 shRNAs, demonstrating that the observed cell depletion was highly specific to the combined loss of STAG1 and STAG2 (**Figure 1D & E**). We reasoned that depletion of both STAG1 and STAG2 in the HSPC model may disrupt cohesin’s essential functions of sister chromatid cohesion during cell division and thereby limit cell survival. (12, 13). Indeed, we observed that STAG1 knockdown induced marked sister chromatid cohesion defects in more than 50% of the *STAG2* null cells (**Figure 1F & G**). Overall, these findings demonstrate the existence of a synthetic lethal interaction between STAG1 and STAG2 in primary human HSPCs.

We then sought to assess whether we could induce synthetic lethality by targeting STAG1 in STAG2-mutated primary AML cells. Primary human AML cells typically survive poorly in culture, and we chose a patient sample that was readily transplantable in immunodeficient mice and that had been propagated as a patient-derived xenograft (PDX) sample to allow assessment *in vivo*. This sample carried a *STAG2* loss-of-function mutation with a variant allele frequency of 89% accompanied with other candidate driver mutations such as IDH2, SRSF2 and NRAS. **(Supplementary table 1)**. We analysed STAG2 protein levels in the PDX sample by Western blotting and found a complete lack of STAG2 expression, in line with the sequencing data **(Figure 2A)**. We successfully transduced bulk mononuclear PDX cells with either scrambled or the two independent STAG shRNAs at frequencies of 30-60% and transplanted into sub-lethally irradiated NOD-scid IL2Rgnull*-3/GM/SF* (NSG-S) mice **(Figure 2B, Supplementary Figure 2A)**. As a reference control, and to assess the effect of STAG1 perturbation on normal HSPCs, we also transduced UCB CD34^+^ cells with the same vectors and assayed the cells both *in vitro* and *in vivo* by transplantation to sub-lethally irradiated NOD.Cg-Prkdc^scid^Il2rg^tm1Wjl^/SzJ (NSG) mice. Importantly, STAG1 knockdown did not appear to negatively impact the engraftment of UCB CD34^+^ cells (**Figure 2C & D)**. Rather, we observed a moderate increase in the fold expansion of CD34^+^ cells *in vitro*, and a robust contribution of transduced cells *in vivo* that was steadily increasing over time, similar to cells transduced with the scramble control (**Supplementary figure 2B & 2C)**. Altogether this indicates that partial perturbation of STAG1 is well tolerated by human HSPCs without a major influence on their repopulation and differentiation properties (**Supplementary figure 2D – 2H)**. By contrast, when we analyzed the bone marrow of NSGS mice transplanted with transduced PDX derived AML cells, we found a near complete depletion of cells transduced with either of the two STAG1 shRNAs, whereas the mice transplanted with scrambled shRNA transduced cells retained a stable transduced KuO^+^ population **(Figure 2E & F)**. This suggests that STAG1 knockdown induces a selective impairment of *STAG2* null AML cells that is sufficient to eliminate them upon transplantation.

**Figure 2.**
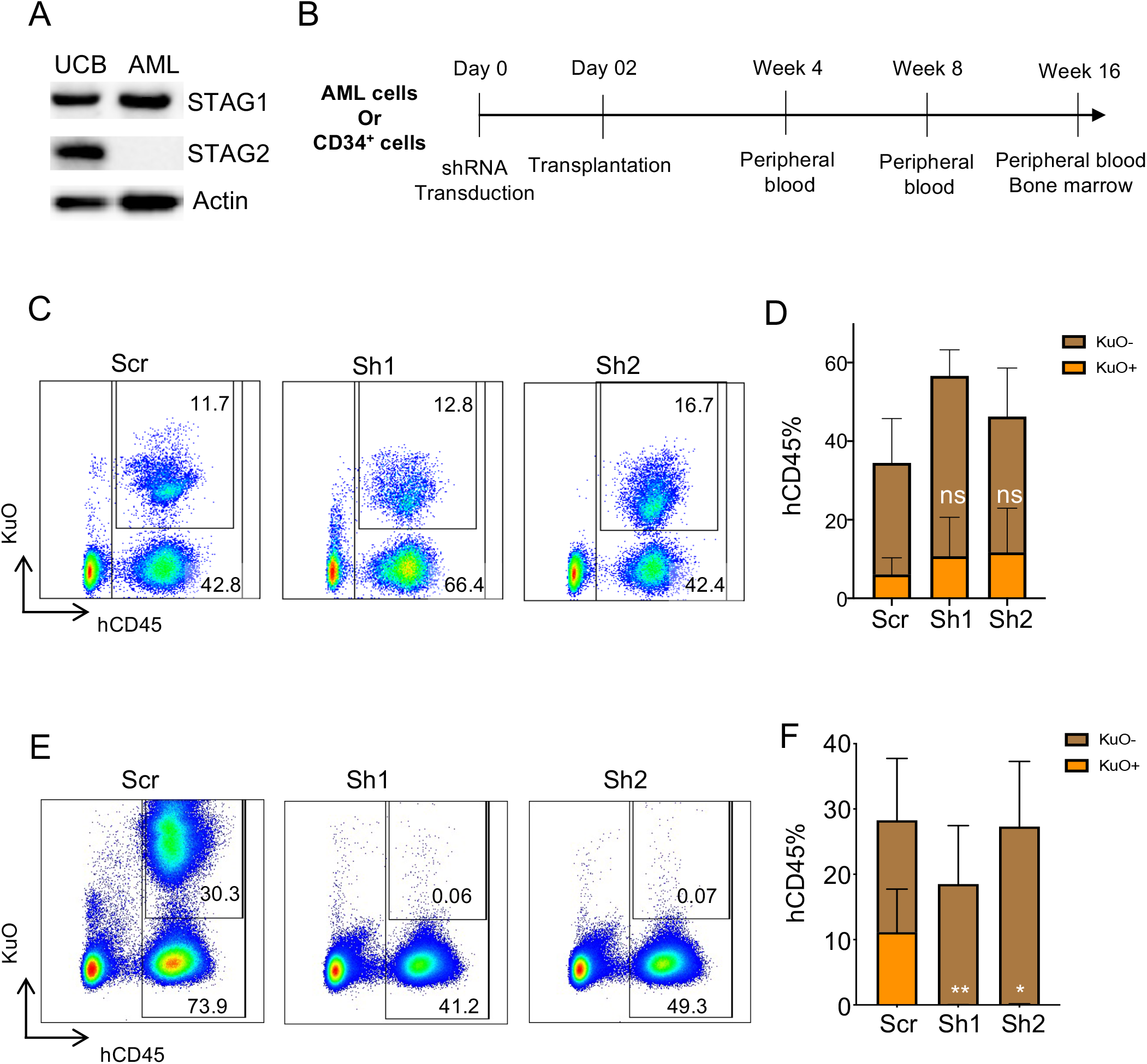
STAG1 knockdown selectively perturbs AML cells. A. STAG1 and STAG2 expression in UCB and *STAG2* null AML cells. B. UCB and AML cells were transduced with scrambled and STAG1 shRNAs and transplanted into sub-lethally irradiated NSG and NSG-S mice respectively. Prior to transplantation AML xenograft cells were transduced *in vitro* with lentiviral particles and transferred to a new plate coated with irradiated OP9 stroma cells. C. FACS plots showing the chimerism of UCB grafts (humanCD45) and frequencies of Kusabira orange positive shRNA transduced cells at NSG bone marrow 16 weeks post transplantation.D. Frequency of hCD45 chimerism and the proportion of transduced cells were quantified for each shRNA (n=5). Kruskal-Wallis test with comparison to the scrambled control. ns-not significant. E. Chimerism of AML grafts (human CD45) and frequencies of Kusabira orange positive shRNA expressing cells in NSG-S bone marrow, analyzed 16 weeks post transplantation. F. Frequency of AML engraftment and the proportion of transduced cells were quantified for each shRNA (n=4). Kruskal-Wallis test with comparison to the scrambled control. p<0.05 - *; p<0.01 - **. Scr-Scrambled shRNA; Sh1-STAG1 shRNA1; Sh2-STAG1 shRNA2.

Taken together, we demonstrate that partial perturbation of STAG1 selectively eliminates primary human HSPCs and AML cells lacking STAG2, while it is well tolerated by normal HSPCs. Developing STAG1 perturbing small molecules would be an ideal way to translate these findings into clinical applications. Our study provides a rationale for exploiting synthetic lethality to develop more specific and targeted therapies for tumors with *STAG2* mutations, demonstrating a first proof-of-concept within the hematopoietic system.

## Acknowledgements

We would like to thank the staff at animal house and FACS core facilities for their excellent support. We also would like to thank Jenny G. Johansson for helping with the AAV vector production. This work was funded to J.L. by grants from the Swedish Research Council, the Swedish Cancer Foundation, the Swedish Pediatric Cancer Foundation, Knut och Alice Wallenbergs Stiftelse and the European Research Council (ERC) under the European Union’s Horizon 2020 research and innovation program (grant agreement No. 648894). A.S. was supported from the Swedish Cancer Foundation, Royal Physiographic Society of Lund and Lady TATA memorial trust. The work was further supported by the HematoLinné and StemTherapy programs at Lund University.

