## Supplementary materials for "Inducing synthetic lethality for selective targeting of acute myeloid leukemia cells harboring *STAG2* mutations"

**Supplementary Methods**

*Cell culture*

CD34^+^ cells were isolated from UCB samples as described previously [1]. Patient derived xenografts for primary AML samples were established as mentioned earlier [2]. UCB and AML samples were collected after informed consent and the study was approved by the regional ethical committee at Skåne University Hospital and Lund University. UCB CD34^+^ cells were cultured and transduced in serum free expansion medium (SFEM) (#09650, Stem cell technologies, Canada) added with stem cell factor (SCF), thrombopoietin (TPO) and FMS-like tyrosine kinase 3 ligand (FLT3L) at final concentration of 100 ng/mL each (#300-07, #300-18, and #300-19, Peprotech, USA). AML PDX cells were cultured and transduced with lentiviral particles overnight in S7 media [3]. After overnight transduction, cells were transferred to a new plate coated with irradiated OP9 stroma cells. Following short hairpin RNA sequences were used, Scrambled- CAACAAGATGAAGAGCACCAA; STAG1 sh1-CTTCAGCCTTTGGTGTTCAAT, STAG1 sh2 -GCCAATGAAAGGTTGGAGTTA.

*Generating STAG2 null cells*

Cas9 RNP complexes containing single guide RNA (UCUGGUCCAAACCGAAUGAA) (Synthego, USA) targeting STAG2 was delivered to UCB HSPCs via electroporation (Harvard Apparatus). To track the successfully edited STAG2 null cells, we delivered a AAV donor template expressing PGK-EGFP flanked with homologous arms spanning 400 bp long each side of the cut site. The AAV donor template was designed according to the detailed protocol by *Tran NT et al., 2020* [4]. AAV 6 was prepared by the AAV Vector Lab (MultiPark, Lund University).

*PCR*

Male samples were genotyped for *SRY* gene while *ATL1* was used as a reference as reported before [5]. To analyze the integration of the PGK-eGFP in *STAG2* locus, primers (Forward TGGGGTAGGCACAGTTTTATCCT and Reverse TGGTTTACATCAGCATATTTTTGACCA) were designed to anneal outside of the homology arms, and the expected sequence of the PCR product was confirmed by Sanger sequencing.

*Cohesion assay*

Sister chromatid cohesion was analyzed as described earlier [6] in the control and sorted KuO^+^ and KuO^+^eGFP^+^ cells, and classified into PCG I to IV according to the severity of defects. The assay was performed in a blinded fashion on fluorescence in situ hybridization (FISH) preparations mounted with DAPI where 8 to 17 cells per each condition was analyzed. Images were captured using a Z2 fluorescence microscope (Zeiss, Germany) and the CytoVision software (Leica, Germany).

*In vivo studies*

UCB CD34^+^ cells were transduced for 48 hours and transplanted into sub-lethally irradiated (3Gy) NSG mice. Peripheral blood samples from tail vein were analyzed for engraftment at 4-, 8- and 16-weeks post transplantation. Bone marrow samples were collected and analyzed at week 16. Animal studies were approved by the local ethical committee at Lund University. AML PDX cells were transduced *in vitro* and cultured for 48 hours and subsequently transplanted into sub-lethally irradiated (3Gy) NSG-S mice and the bone marrow was analyzed 16 weeks post transplantation

*Flow cytometry*

LSR Fortessa and FACS Canto II (Becton Dickinson) were used to analyze the samples. Cell sorting was performed in FACS Aria III (BD). FACS antibodies: CD34-FITC (#343604), EPCR-APC (#351906), CD33-FITC (#303304), CD38-PE-Cy7 (#303516), CD45-APC (#304012) and CD34-PE-Cy7 (#343516) from BioLegend, UK; CD19-BV605 (#562653) from BD; CD3-PE-Cy7 (#25-0038-42) from eBioscience.

*Western Blotting*

STAG2 edited eGFP positive population was sorted and collected for Western analysis. shRNA transduced CD34^+^ cells were collected 48hours post-transduction. Cells were washed 3 times with PBS and lysed with 2X Laemmli buffer at 95^o^C for 5 minutes. Western blotting was performed with NuPAGE system following manufacturer’s instructions (ThermoFisher, Manual part no. IM-1001). Primary antibodies: STAG1 (# ab4457) from abcam; STAG2 (#4239) from Cell Signaling Technology and Actin (#612656) from Becton Dickinson.

**Supplementary figure legends**

**Supplementary Figure 1.** A. Construction of the eGFP knock in construct and validation of the integration in the STAG2 locus with PCR. B. Integration of eGFP construct was analyzed with PCR with the following primers, Primer 1; TGGGGTAGGCACAGTTTTATCCT and Primer 2; TGGTTTACATCAGCATATTTTTGACCA. C. UCB CD34^+^ cells were transduced with STAG1 shRNAs for 3 days and the knockdown efficiency was quantified with qPCR and D. with Western blotting.

**Supplementary Figure 2.** A. Transduction frequencies of scrambled and STAG1 shRNAs in AML PDX, two days post transduction, quantified from 3 independent replicates. B. UCB CD34^+^ cells were transduced with STAG1 shRNAs and fold expansion of CD34 cell numbers were quantified over four weeks of *in vitro* culture. C. UCB CD34^+^ cells were transduced with STAG1 shRNAs and transplanted in NSG mice. Engraftment of transduced Kusabira orange positive cells in the peripheral blood at week 4, 8 and 16. D. CD19 B-cell and CD33 myeloid lineage contribution of shRNA transduced cells in bone marrow at week 16. E. B-, Myeloid and T- cell lineage frequencies were quantified for each shRNA. F. Frequency of CD34^+^ and CD34^+^CD38^low^ cells in observed in the bone marrow of NSG mice transplanted with shRNA transduced UCB CD34^+^ cells. Quantification of the frequency of G. CD34^+^ and H. CD34^+^CD38^low^ cells for each shRNA.

**Supplementary table 1.** Next generation sequencing analysis-based frequencies of the somatic mutations in AML patient-derived xenograft (PDX) cells.
