## Supplementary figures for "Inducing synthetic lethality for selective targeting of acute myeloid leukemia cells harboring *STAG2* mutations"

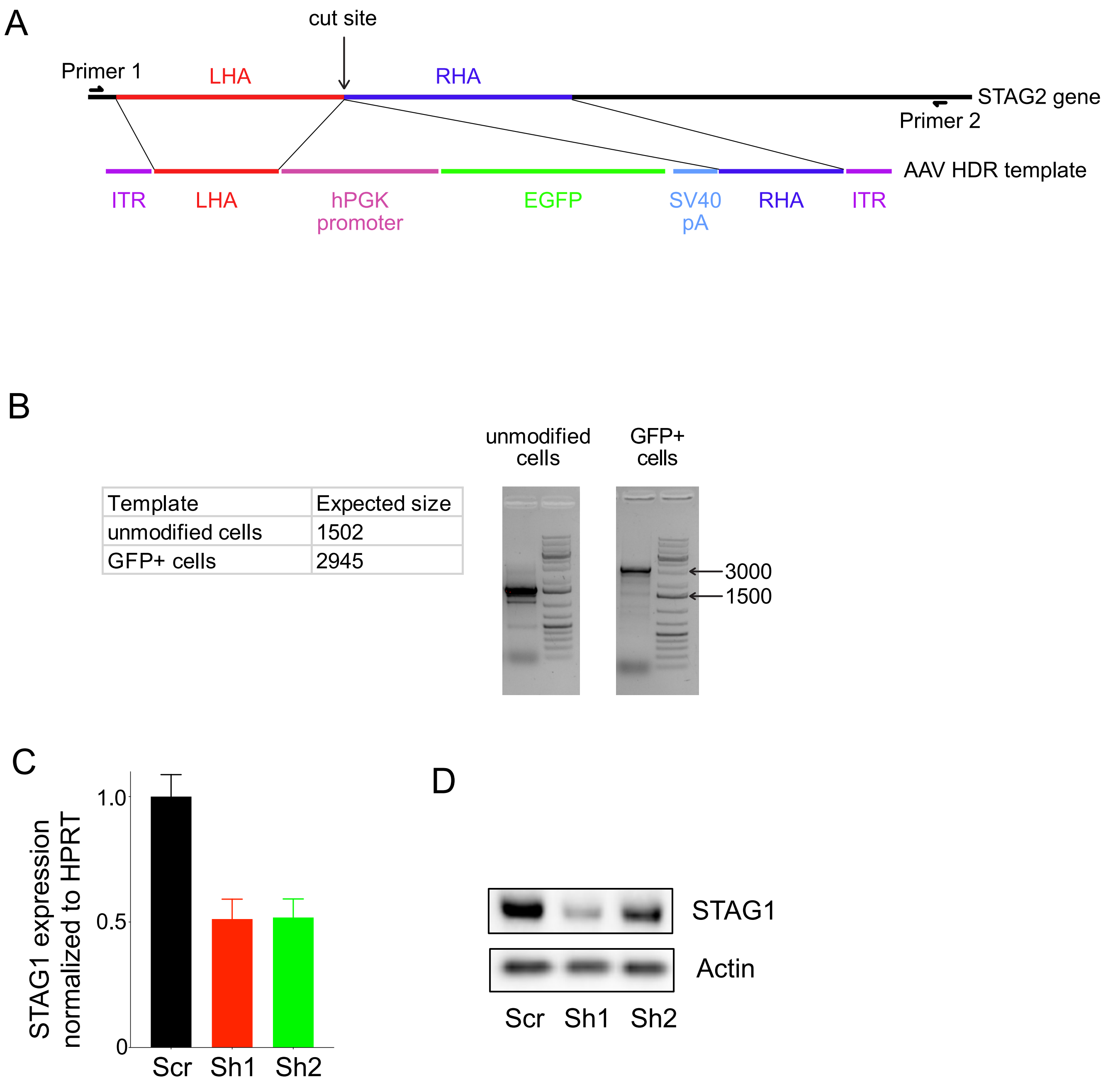


**Supplementary Figure 1**

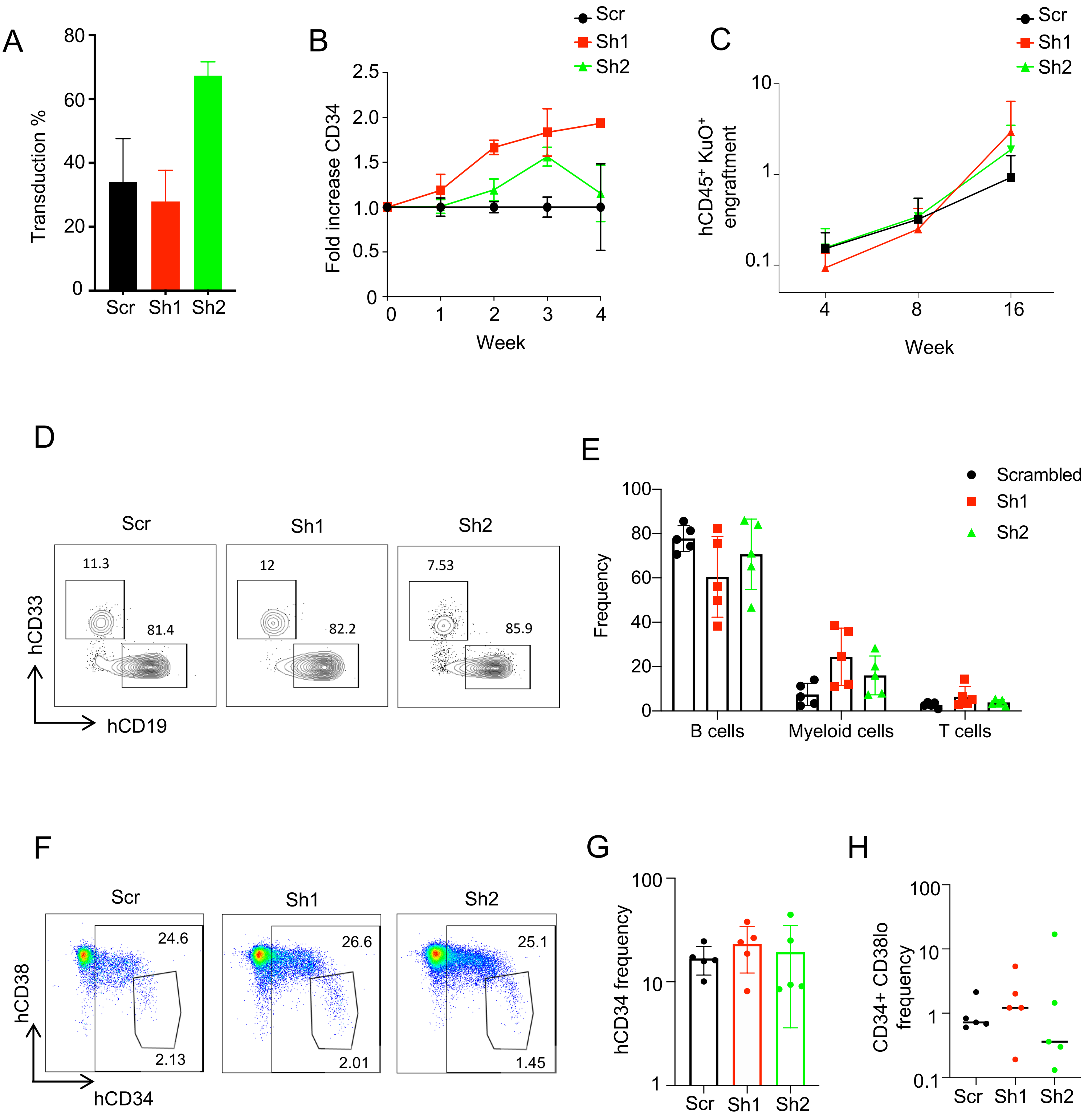


**Supplementary Figure 2**

| Gene | Chromosome | Start position | End position | Reference sequence | Alternative sequence | Sequencing depth | Variant allele frequency |
|---|---|---|---|---|---|---|---|
| *STAG2* | chrX | 123220481 | 123220486 | TTCTTT | TGAAAGCT | 37 | 0.89 |
| *GRIA4* | chr11 | 105758236 | 105758236 | T | C | 76 | 0.64 |
| *MPP4* | chr2 | 202547667 | 202547667 | A | C | 28 | 0.64 |
| *EDAR* | chr2 | 109546594 | 109546594 | C | T | 24 | 0.63 |
| *HNF1A* | chr12 | 121432080 | 121432080 | C | T | 178 | 0.55 |
| *TUBB3* | chr16 | 89999064 | 89999064 | G | A | 79 | 0.48 |
| *NRAS* | chr1 | 115258747 | 115258747 | C | T | 172 | 0.47 |
| *SERPINB4* | chr18 | 61310786 | 61310786 | G | A | 129 | 0.46 |
| *PROX1* | chr1 | 214170676 | 214170676 | G | A | 79 | 0.46 |
| *DRD5* | chr4 | 9784632 | 9784632 | G | A | 66 | 0.45 |
| *IDH2* | chr15 | 90631934 | 90631934 | C | T | 171 | 0.45 |
| *TTN* | chr2 | 179436904 | 179436904 | A | G | 78 | 0.45 |
| *ANXA2P2* | chr9 | 33625306 | 33625306 | T | C | 45 | 0.44 |
| *SRSF2* | chr17 | 74732959 | 74732959 | G | T | 71 | 0.44 |
| *SH2B3* | chr12 | 111885251 | 111885251 | T | C | 180 | 0.43 |
| *SARS* | chr1 | 109779189 | 109779189 | G | C | 27 | 0.41 |
| *TGFBR3* | chr1 | 92149368 | 92149368 | C | T | 105 | 0.40 |
| *TGM4* | chr3 | 44952468 | 44952468 | C | T | 49 | 0.39 |
| *MMP13* | chr11 | 102820988 | 102820988 | G | C | 122 | 0.39 |
| *TANGO6* | chr16 | 68936253 | 68936254 | TT | TCACCTTATGGGGCTCTAA | 19 | 0.16 |

**Supplementary table 1**
